# The RNA from *Pseudomonas aeruginosa* impairs neutrophil responses favoring bacterial survival

**DOI:** 10.1101/2024.01.10.574964

**Authors:** José R Pittaluga, Federico Birnberg-Weiss, Agustina Serafino, Joselyn Castro, Luis A Castillo, Daiana Martire-Greco, Paula Barrionuevo, Gabriela C Fernández, Verónica I Landoni

## Abstract

Epithelial and endothelial cells are essential in the modulation of innate immune responses in the lung, including the arrival of neutrophils (PMN), which are crucial cells for the antibacterial host defense. These cells are exposed to prokaryotic RNA (pRNA) during bacterial infections and pRNA has been shown to promote or attenuate the inflammatory response on different immune cells. *Pseudomonas aeruginosa* (PAE) can cause severe pneumonia and has several immune-evading mechanisms. The aim of this study was to determine the effects of the RNA from PAE (RNA_PAE_) on lung epithelial, endothelial cells and PMN, and its impact on bacterial elimination. For this purpose, we purified total RNA_PAE_, and used it as a stimulus to evaluate different functions on Calu-6, HMEC-1 and isolated human PMN. We found that RNA_PAE_ neither induced a pro-inflammatory response on Calu-6 or HMEC-1, as measured by ICAM-1 surface expression, or IL-6 and IL-8 secretion. Also, RNA_PAE_ failed to activate PMN, as measured by forward-scatter (FSC) increase, CD11b surface expression, chemotaxis and IL-8 secretion. Pre-stimulation with RNA_PAE_ diminished CD11b surface expression, chemotaxis and microbicidal activity when PMN were challenged with live bacteria. Moreover, we found that phagocytosis was affected in the presence of RNA_PAE_. Fragments of short RNA (<200 bp) were responsible for the PMN microbicidal attenuation during bacterial elimination. In conclusion, our results indicated that short fragments of RNA_PAE_ diminished the immune response on PMN even in the presence of live bacteria.

**AUTHOR SUMMARY:** *Pseudomonas aeruginosa* (PAE) pneumonia constitutes a major problem for human health. Therapies are frequently inefficient due to immune evasion mechanisms of PAE. Therefore, it is imperative to understand the relationship between PAE (or its components) with the immune system to improve therapeutic strategies.

Since some bacterial RNA are immunosuppressive, our hypothesis was that the RNA from PAE (RNA_PAE_) might negatively modulate the immune response in a lung infection. We investigated the effects of the RNA_PAE_ on lung epithelial, and microvascular endothelial cells, central cells that respond to PAE early during infection, and on neutrophils (PMN), the first immune cell that arrives at the site of infection.

We found that RNA_PAE_ failed to induce any response on pulmonary epithelium, endothelium, or PMN. Moreover, RNA_PAE_-treated PMN showed reduced migration, activation, and bactericidal response against live bacteria. Exploring deeper into this phenomenon, we found that increased bacterial survival was due to a lower phagocytic capacity of RNA_PAE_-treated PMN.

Our results indicate that RNA_PAE_ may act as another evasion strategy to favor PAÉs survival in a pulmonary infection. Understanding the mechanisms by which PAE reduces the response of cells that participate in pulmonary immunity is crucial for planning interventions that may benefit infected patients.

## INTRODUCTION

Nosocomial infections continue to be a major threat for hospitalized patients and a burden to healthcare institutions, causing great economic losses to the health system. Severe pneumonia acquired in healthcare centers are caused mainly due to Gram-negative organisms (1,2), such as *Pseudomonas aeruginosa* (PAE) (3,4), and members of the Enterobacteriaceae family, such as *Escherichia coli* (ECO) (1,5,6). In general, these are opportunistic microorganisms that can colonize both healthcare personnel and patients, especially those admitted to intensive care units (ICU), and/or immune compromised.

These bacteria are continuously regulating and acquiring genes that code for mechanisms of antibiotic resistance (7), and are listed by the World Health Organization (WHO) as bacteria of priority research (8). Resistance to antibiotics is an increasingly concerning problem, and elucidating the pathophysiology of bacterial infections is essential to establish the basis and tools that could effectively solve this global problem.

The lung barrier, formed by epithelial and endothelial cells, is essential in the modulation of the innate immune response induced in the lung (9,10). The arrival of immune cells to infected lungs strongly depends on epithelial and endothelial cell activation. Secretion of inflammatory and chemoattractant cytokines along with adhesion molecule expression on cell membranes are required to favor immune cell adhesion and transmigration. Neutrophils (PMN) are the first line of antibacterial host defense and are activated through cytokine released at the site of infection. This promotes PMN attraction, adherence, and transmigration across the endothelium (11). Once PMN reach the site of infection, their role in bacterial clearance relies on ingestion, and intracellular and extracellular killing of bacteria.

Bacterial prokaryotic RNA (pRNA) has been found at the site of infection (12) and also in blood (13). pRNA can be secreted by bacteria as a mechanism of defense, associated with biofilm formation or inside bacterial-released microvesicles (14,15). In particular, it has been reported that, as an adaptation to the environment, PAE releases its nucleic acids, which acts in its metabolic regulation, adaptation to stress, virulence and biofilm formation (16). Moreover, it is also considered that pRNA in the extracellular matrix can also be present due to bacterial killing by immune cells (17–19). In this sense, a dead bacteria, when eliminated by the immune system, will release large amounts of RNA (10 times more than DNA) (17,18). Therefore, mucosal cells and nearby tissues are regularly exposed to the RNA of microorganisms that have been killed. Released pRNA impacts host cells and this can directly modulate the host response (20–22). It is beginning to be revealed that “riboregulation” may be central to the pathogenic mechanisms of certain pathogens (23,24).

There is evidence about the immunomodulatory capacity of pRNA from some bacteria, but the effects of these RNAs are variable depending on the type of bacteria and the target cell. While some studies have shown that pRNA can induce an inflammatory response in different immune cells (18,25,26), other studies have documented that pRNA mediates immune evasion favoring bacterial survival and colonization (27,28).

It is known that PAE possess multiples mechanisms that alters the immune response (29) and some virulence factors from PAE manipulate the inflammatory response to avoid killing (30). Therefore, our hypothesis is that RNA from PAE may modulate the lung barrier and PMN immune responses favoring bacterial survival.

Considering this, the aim of this study was to determine the effects of RNA from PAE (RNA_PAE_) on lung epithelial, endothelial cells, and PMN, as well as the impact of this response on bacterial elimination.

## RESULTS

### RNA from *P. aeruginosa* failed to induce an inflammatory response on endothelial and lung epithelial cell

The induction of a response in lung epithelial cells as well as endothelial cells close to an infection focus is crucial to establish an effective immune response against bacteria, specially modulating the recruitment and transmigration of immune cells to the lung. The presence and leakage of pRNA into the nearby vasculature could influence this response. Therefore, we first evaluate the effects of RNA_PAE_ on endothelial and epithelial cell activation and cytokine secretion.

Human lung epithelial cells (Calu-6) or human microvascular endothelial cells (HMEC-1) were stimulated for 18 h with RNA_PAE_ or RNA_ECO_ for comparison. ICAM-1 expression was determined by flow cytometry, and secreted cytokines on cell culture supernatants were measured using ELISA kits.

Figure 1A shows that stimulation of Calu-6 with RNA_ECO_ induced an increase in ICAM-1 expression compared to unstimulated (Control) cells. Surprisingly, RNA_PAE_ failed to induce ICAM-1 expression showing no differences compared to control cells.

**Fig 1.**
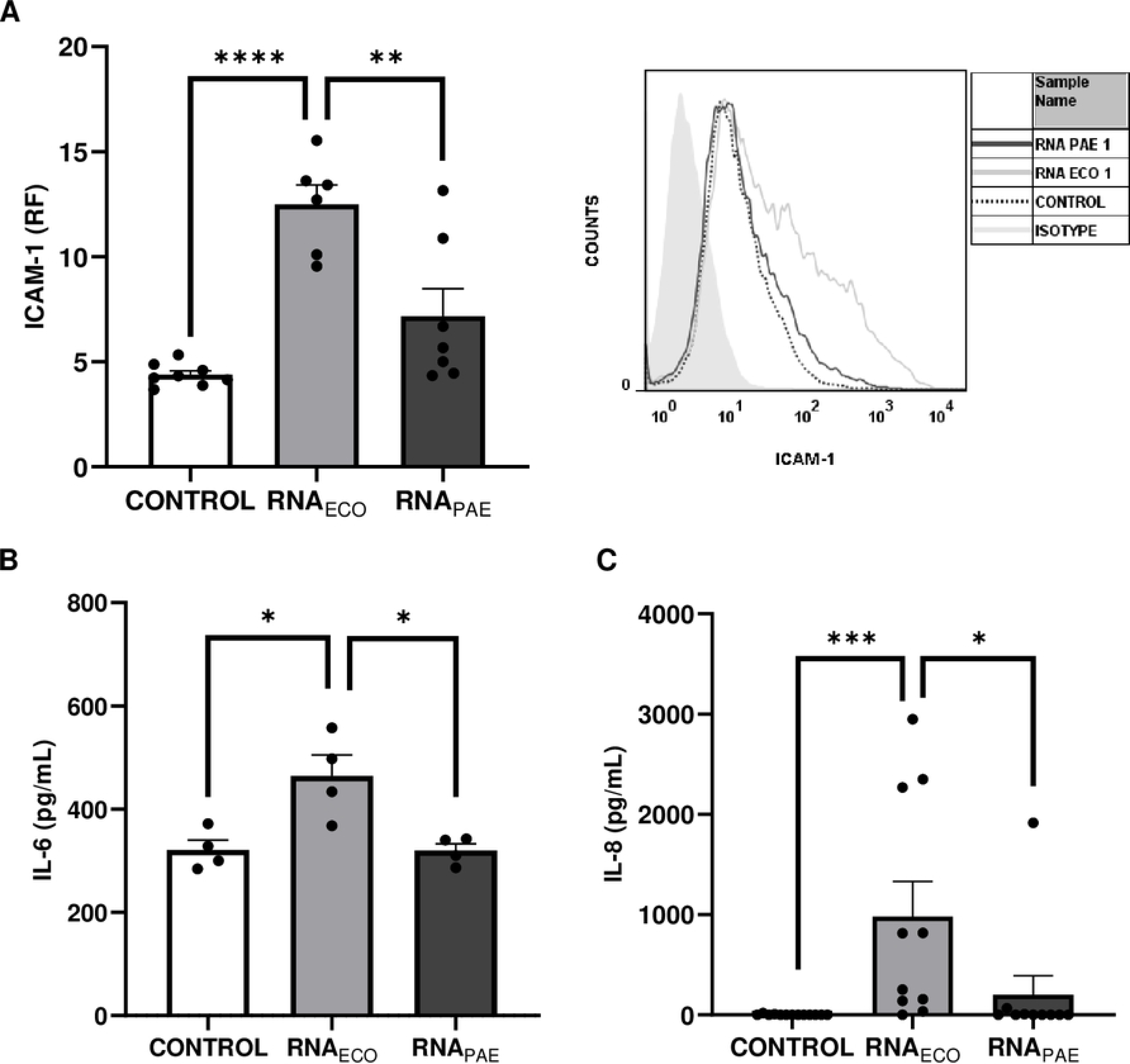
Activation of epithelial lung cells induced by pRNA. A Calu-6 cell line was unstimulated (CONTROL) or stimulated with 1 µg/mL RNA (RNA_PAE_ or RNA_ECO_) for 18 h and then cells and conditioned medium (CM) were collected. **(A)** ICAM-1 expression on cell surface by flow cytometry was expressed as the relative fluorescence (RF) of the mean fluorescence intensity (MFI) of treated samples relative to the isotype control. An histogram plot of one representative experiment is presented (right). **(B)** IL-6 and IL-8 secretion were determined in the CM by ELISA. **(B)** IL-6 concentration. **(C)** IL-8 concentration. Each point represents an independent experiment. Results are expressed as mean ± SEM (*p<0.05, **p<0.01, ***p<0.001, ****p<0.0001).

When the presence of secreted IL-6 or IL-8 chemokine were determined in conditioned medium (CM), significantly increased amounts of both cytokines were found when RNA_ECO_ was used as a stimulus (Figure 1B **and 1C**). Again, IL-6 or IL-8 secreted levels with RNA_PAE_ were not different compared to control cells.

We next determined the effect of RNA_PAE_ and RNA_ECO_ on HMEC-1 response. As shown in Figure 2A, stimulation of HMEC-1 with RNA_ECO_ significantly increased ICAM-1 expression compared to untreated (control) cells, as well as induced higher amounts of secreted IL-6 (Figure 2B) and IL-8 (Figure 2C). As with epithelial cells, RNA_PAE_ was not able to activate endothelial cells (ICAM-1 expression), and IL-6 or IL-8 secretion were not statistically different compared to control cells.

**Fig 2.**
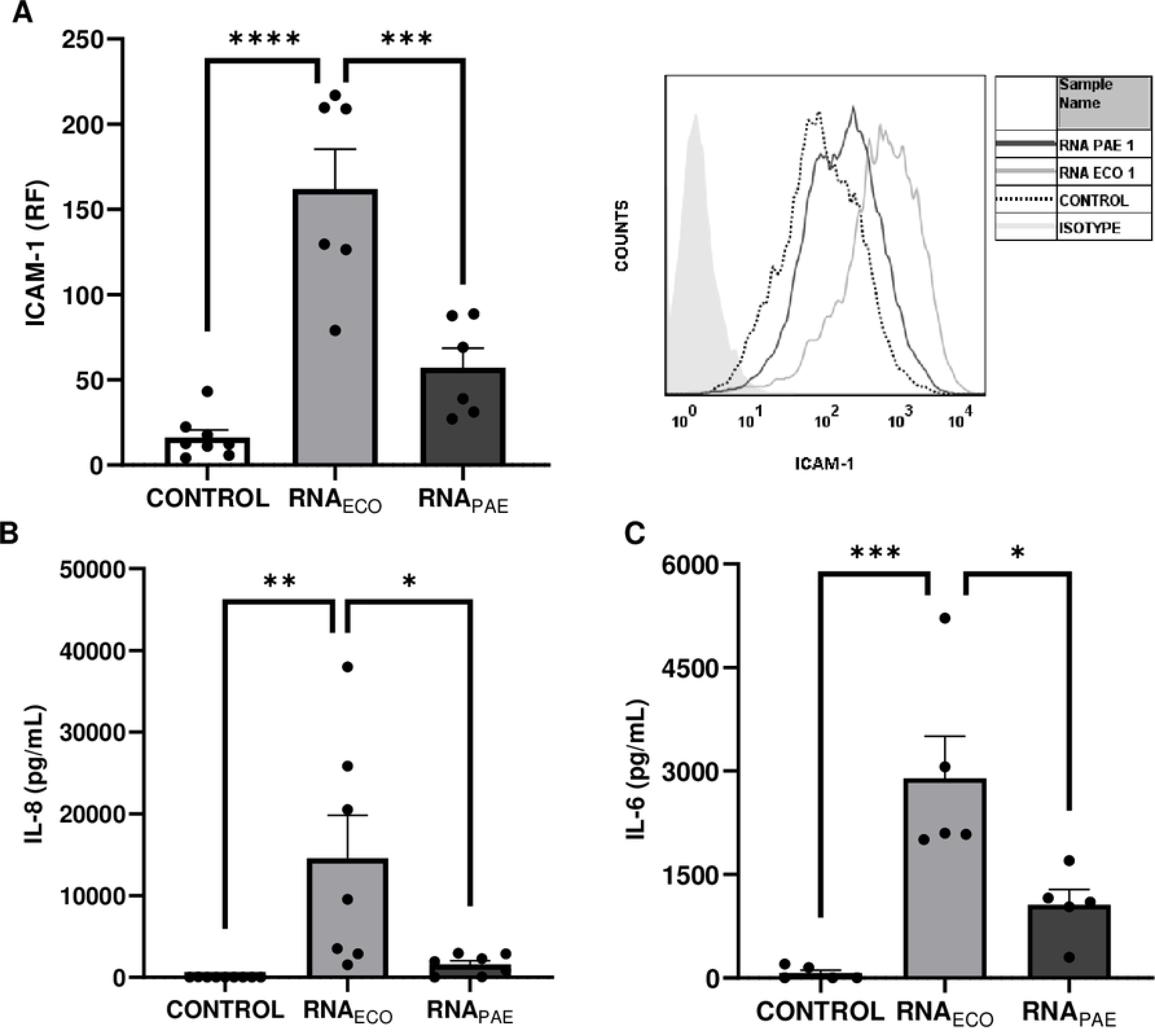
Activation of microvascular endothelial cells induced by pRNA. HMEC-1 cells were unstimulated (CONTROL) or stimulated with 1 µg/mL RNA (RNA_PAE_ or RNA_ECO_) for 18 h and then cells and conditioned medium (CM) were collected. **(A)** ICAM-1 expression on cell surface by flow cytometry was expressed as the relative fluorescence (RF) of the mean fluorescence intensity (MFI) of treated samples relative to isotype control. A histogram plot of one representative experiment is depicted (right). **(B)** IL-6 and IL-8 secretion were determined in the CM by ELISA. **(B)** IL-6 concentration. **(C)** IL-8 concentration. Each point represents an independent experiment. Results are expressed as mean ± SEM (*p<0.05, **p<0.01, ***p<0.001, ****p<0.0001).

These results demonstrate that RNA_PAE_ failed to induce a pro-inflammatory response on epithelial or endothelial cells in contrast to RNA_ECO_.

A five times higher dose of RNA_PAE_ was also used to stimulate cells, in order to corroborate that the non-stimulatory effects could be ascribed to a low stimulus dose. As shown in **Supplementary** Figure 1, a higher dose of RNA_PAE_ also failed to activate epithelial or endothelial cells. Also, cytotoxicity assays were performed incubating cells overnight with the different pRNAs, and no significant death was registered among treatments (**Supplementary** Figure 2A and 2B).

### RNA from *P. aeruginosa* failed to induce an inflammatory response on PMN

As many immune cells, PMN have receptors to detect RNA (31), and considering that pRNA has been detected in the bloodstream (13), it is likely that bacterial RNA nearby the site of infection could directly contact PMN. Since PMN are the first cells to reach the infectious site and bacterial pneumonia are characterized by a strong neutrophilic response (32), we wanted to determine the effects of pRNA on PMN functions.

Firstly, to rule out any toxic effect on PMN caused by pRNA, cytotoxicity assays were performed incubating pRNA for 4 h with isolated PMN, and no significant cell death was observed with RNA_PAE_ or RNA_ECO_ (**Supplementary** Figure 2C).

Then, PMN from healthy donors were stimulated with 1 µg/ml of pRNA for 30 min. Parameters of activation (CD11b and changes in cell size) were analyzed by flow cytometry. As shown in Figure 3, RNA_PAE_ failed to induce an up-regulation in CD11b expression (Figure 3A) or any change in cell size measured as the % of cells with high forward scatter (FSC) (Figure 3B). Even at higher concentrations, RNA_PAE_ was not able to induce PMN activation (**Supplementary** Figure 3A).

**Fig 3.**
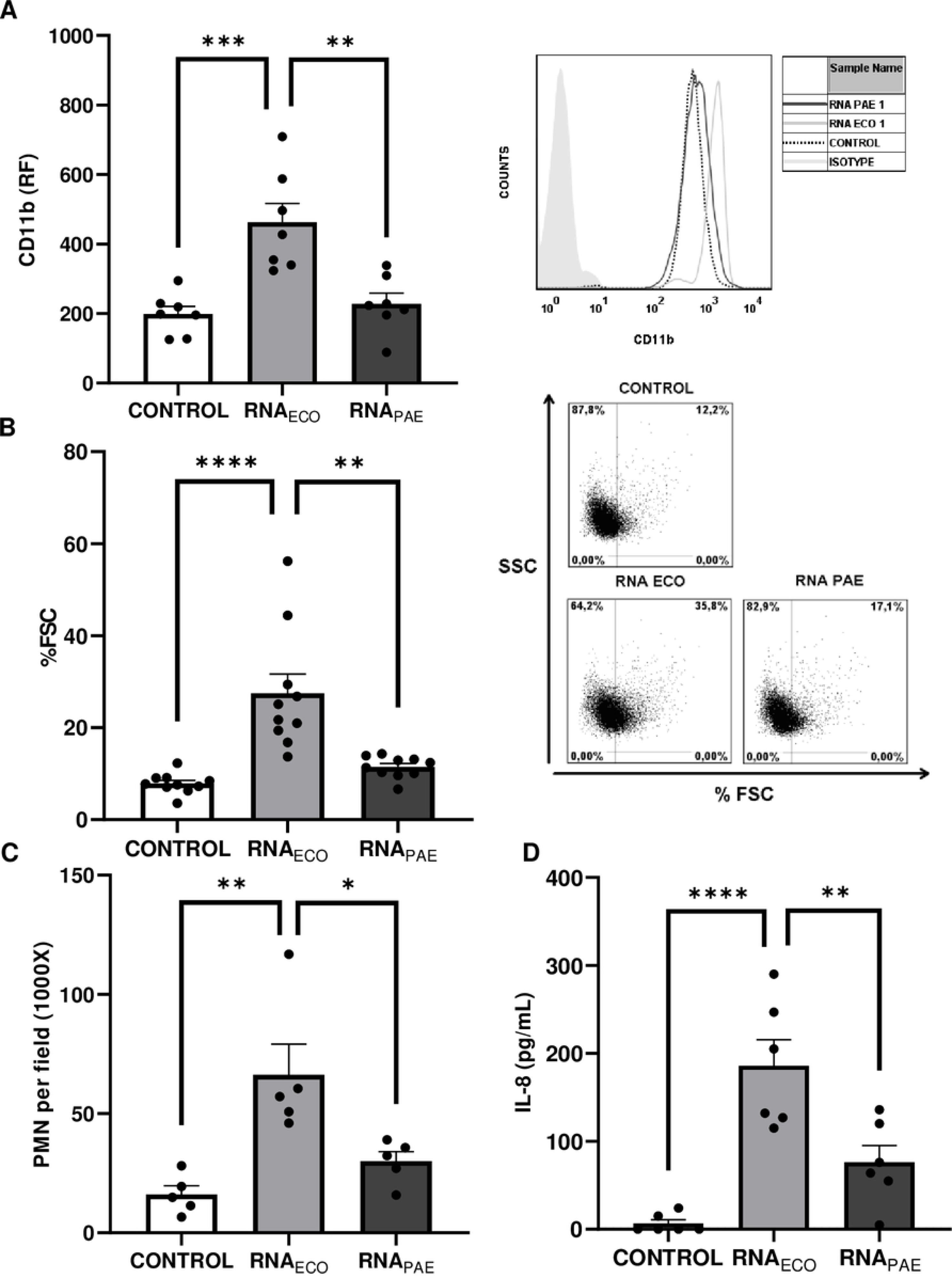
Activation responses on neutrophils (PMN) induced by pRNA. PMN were unstimulated (CONTROL) or stimulated with 1 µg/mL RNA (RNA_PAE_ or RNA_ECO_) for 30 min and CD11b expression (**A)** and changes in the cell size (% FSC) **(B)** were measured by flow cytometry**. (C)** Chemotaxis induced by pRNA was determined using a Boyden chamber**. (D)** IL-8 secretion was measured by ELISA in the supernatant of treated-PMN after 4 hours of stimulation. **(A)** CD11b was expressed as the relative fluorescence (RF) of the mean fluorescence intensity (MFI) of treated samples relative to the isotype control. An histogram plot of one representative experiment is presented (right). **(B)** % FSC of PMN. Representative dot-plots are depicted in the right. **(C)** Number of migrated PMN *per* field (1000X) is shown. **(D)** IL-8 concentration. Each point represents an independent PMN donor. Results are expressed as mean ± SEM (*p<0.05, **p<0.01, ***p<0.001, ****p<0.0001).

Next, we investigated chemotactic properties of pRNA (1 µg/ml) using a Boyden chamber. As shown in Figure 3C, RNA_ECO_ induced chemotaxis of PMN compared to control. However, RNA_PAE_ failed to induce chemotaxis of PMN.

Additionally, secretion of IL-8 was evaluated after 4 h of stimulation with pRNA (1 µg/ml). Figure 3D shows that RNA_ECO_ was able to induce IL-8 secretion compared to unstimulated PMN (control), whereas RNA_PAE_ was not able to induce this cytokine even at a higher concentration (**Supplementary** Figure 3B).

In summary, data presented here demonstrates that RNA_PAE_ was not able to induce a pro-inflammatory response on PMN compared to RNA_ECO_.

### RNA from *P. aeruginosa* failed to induce secretion of PMN-activating factors from epithelial or endothelial cells

Up to this point, we have studied the direct effects of RNA_PAE_ and RNA_ECO_ on PMN, lung epithelial and endothelial cells. However, soluble factors released by endothelial and epithelial cells at the infection site could affect circulating immune cells, particularly influencing their arrival and extravasation. Considering this, we next evaluated whether secreted factors induced by pRNA-stimulated Calu-6 or HMEC-1 could modulate PMN responses.

Calu-6 or HMEC-1 were treated with pRNA (1 µg/ml) for 4 h, then washed twice to remove all pRNA and incubated for another 18 h. These RNA-free CM from treated cells (CM-RNA_ECO_ or CM-RNA_PAE_) were used to stimulate PMN.

Figure 4A shows an increase in CD11b expression on PMN stimulated with CM-RNA_ECO_ from Calu-6 (Figure 4A) or HMEC-1 (Figure 4B) cells compared to PMN stimulated with CM from unstimulated cells (CM-Control). In line with this, and according to the levels of secreted IL-8 described in Figure 1C and 2C, chemotaxis of PMN induced by CM-RNA_ECO_ from both Calu-6 and HMEC-1 was also statistically higher than the one observed using CM-Control (Figure 4C and **4D**).

**Fig 4.**
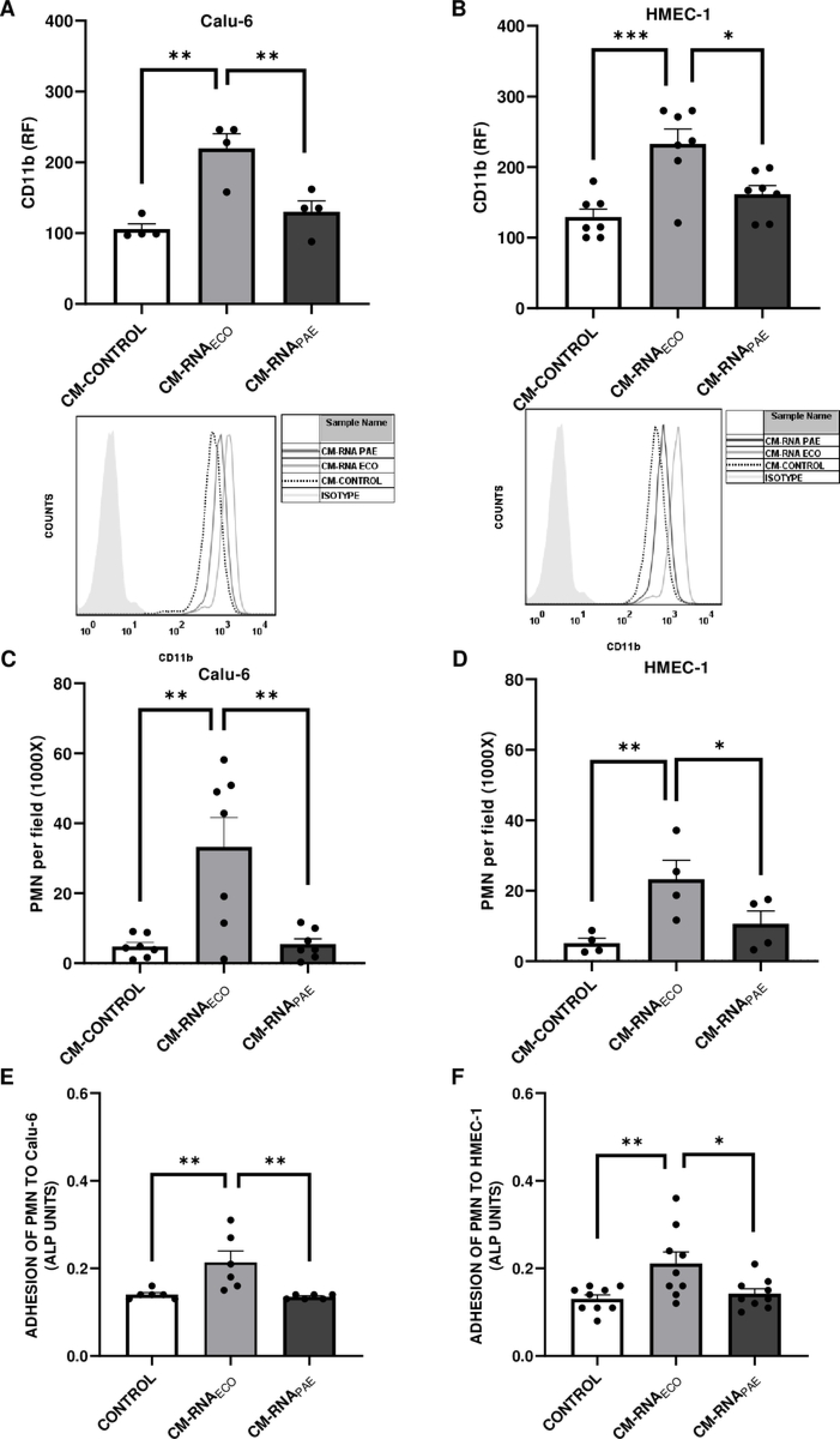
Activation responses of PMN induced by conditioned medium (CM) from epithelial lung or endothelial cells treated with pRNA. Calu-6 or HMEC-1 were treated with 1 µg/mL RNA (RNA_PAE_ or RNA_ECO_) for 4 h, then washed to remove the stimulus and incubated for another 18 h. RNA-free conditioned medium (CM) from Calu-6 **(A and C)** or HMEC-1 **(B and D)** were used to measure modulation of PMN response. Chemotaxis induced by CM was determined using a Boyden chamber **(C, D). (E, F)** PMN adhesion to Calu-6 (E) or HMEC-1 (F) was measured using alkaline phosphatase (ALP) activity, and results were expressed as ALP units **(A, B)** CD11b expression was measured by flow cytometry. CD11b was expressed as the relative fluorescence (RF) of the mean fluorescence intensity (MFI) of treated samples relative to isotype control. A histogram plot of one representative experiment is depicted (below). **(C, D)** Number of migrated PMN *per* field (1000X). **(E, F)** ALP units. Each point represents an independent PMN donor. Results are expressed as mean ± SEM (*p<0.05, **p<0.01, ***p<0.001).

As expected, given the lack of direct effect of RNA_PAE_ on Calu-6 or HMEC-1 cells shown in Figures 1 and 2, when PMN were stimulated with CM-RNA_PAE_ from Calu-6 or HMEC-1, no modulation of CD11b (Figure 4A and **4B**) or chemotaxis (Figure 4C and **4D**) were observed

Finally, adhesion of PMN to epithelial or endothelial cells was also evaluated. Cell lines (Calu-6 or HMEC-1) were not treated (Control) or treated with RNA_ECO_ or RNA_PAE_ (1 µg/ml) for 4 h and then fresh medium was replaced and left for 18 h before PMN were seeded. After 90 min, non-adhered PMN were vigorous washed and adhered PMN were determined by measuring alkaline phosphatase (ALP) activity (see Material and methods). Results depicted in Figure 4E **and 4F** indicate that only RNA_ECO_-treated epithelial and endothelial cells were able to induce PMN adhesion.

These results indicate that factors secreted by Calu-6 or HMEC-1 after RNA_ECO_ stimulation are able to induce activation of PMN favoring their adhesion to cells, while RNA_PAE_ failed to induce secretion of PMN-activating factors.

### RNA from *P. aeruginosa* reduces PMN responses triggered by live bacteria

According to our results, we wondered whether RNA_PAE_ resulted simply not stimulatory or it could have suppressing effects on PMN by modulating their response upon bacterial encounter at the infectious focus in the lung.

Therefore, we evaluated PMN responses against live bacteria in the presence or absence of RNA_PAE_. PMN were stimulated with RNA_PAE_ for 20 min before bacterial challenge (MOI 1). Figure 5A shows that the increase in CD11b expression caused by live PAE was statistically minor when PMN were pre-stimulated with RNA_PAE_. On the contrary, a slightly but statistically significant increase on CD11b expression was registered when PMN were pre-stimulated with RNA_ECO_ and then challenged with live ECO (Figure 5B).

**Fig 5.**
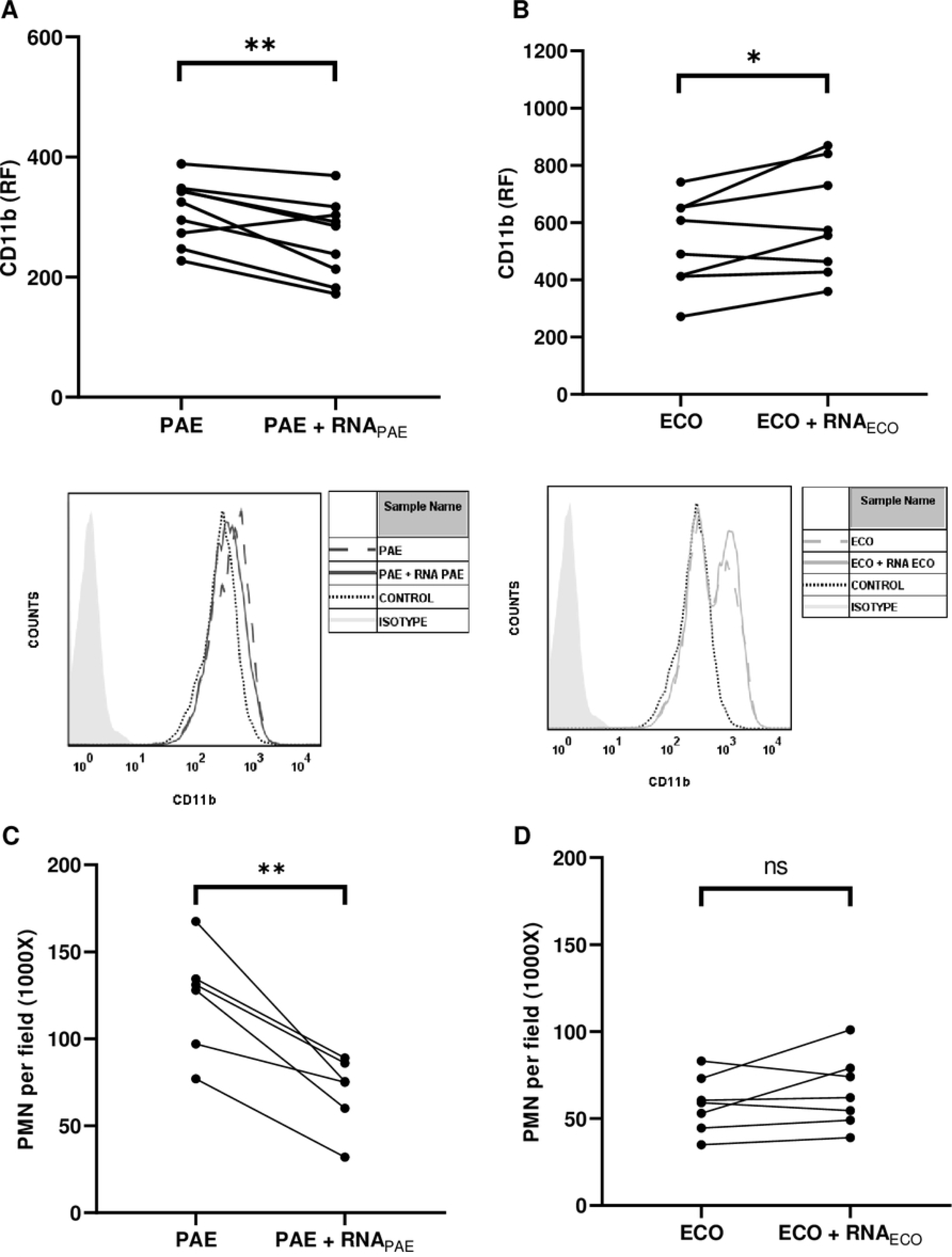
Activation responses on neutrophils (PMN) induced by bacteria in the presence of pRNA. PMN were stimulated with 1 µg/mL RNA (RNA_PAE_ or RNA_ECO_) 20 min before bacterial challenge (MOI 1) for 30 min. **(A)** CD11b expression by PAE after pre-incubation with RNA_PAE_ was measured by flow cytometry **(B)** CD11b expression induced by ECO after not treated or treated with RNA_ECO_**. (C and D)** Migration induced by bacteria on PMN treated or not with pRNA was determined using a Boyden chamber. **(A and B)** CD11b was expressed as the relative fluorescence (RF) of the mean fluorescence intensity (MFI) of treated samples relative to the isotype control. An histogram plot of one representative experiment is presented (below). **(C** and **D)** Number of migrated PMN *per* field (1000X). Each point represents an independent PMN donor. Results are expressed as the mean ± SEM (*p<0.05, **p<0.01).

Moreover, using a Boyden chamber chemotaxis assay towards live bacteria, we found that the number of migrated PMN in the presence of live PAE was significantly reduced by pre-stimulation with RNA_PAE_ (Figure 5C), while chemotaxis of PMN induced by live ECO was not altered by pre-incubation with RNA_ECO_ (Figure 5D).

These results indicate that RNA_PAE_ was inhibitory of PMN functions and its presence reduced the ability of PMN to become activated and migrate towards bacteria.

### RNA from *P. aeruginosa* decreases PMN microbicidal ability

In order to determine whether RNA_PAE_ reduction effects on PMN responses could result in an advantage for bacterial survival, we performed a killing assay. For this purpose, PMN were left unstimulated or were incubated with pRNA for 20 min, and then were challenged with bacteria for 1 h. Cultures were lysed and seeded on TSA plates for determining the number of Colony Forming Units (CFU).

As shown in Figure 6A the number of CFU of PAE was higher when PMN were pre-stimulated with RNA_PAE_ compared to non stimulated PMN. On the other hand, the number of CFU of ECO was not different in the presence of RNA_ECO_ (Figure 6B).

**Fig 6.**
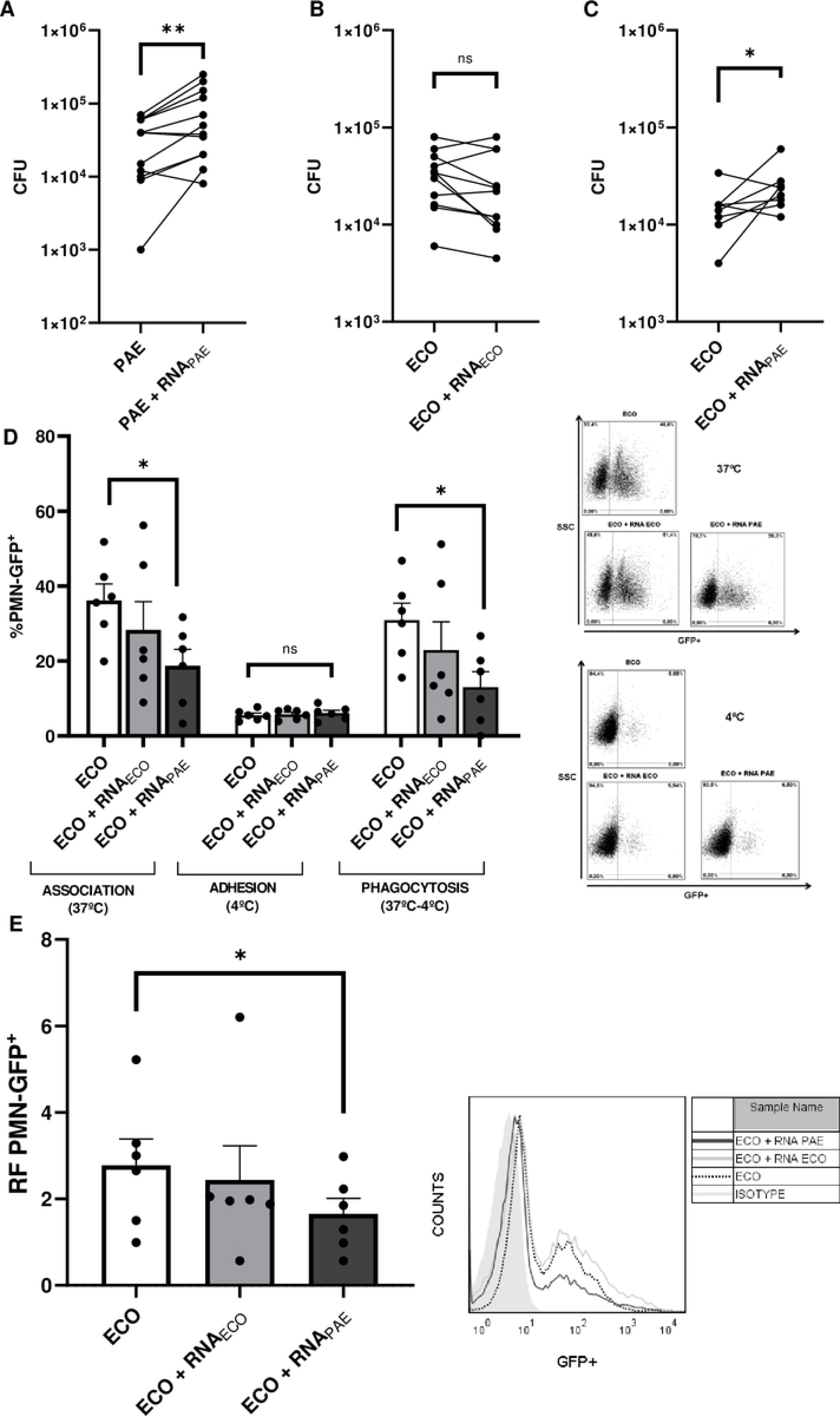
Killing response of PMN stimulated with pRNA. PMN were challenged with PAE or ECO for 1 h (MOI 1) in the presence or absence of 1 µg/mL RNA (RNA_PAE_ or RNA_ECO_) and the remaining viable colony forming units (CFU) were evaluated using TSA plates. **(A)** CFU of PAE after co-incubation of PMN treated or not with RNA_PAE_. **(B)** CFU of ECO after co-incubation of PMN treated or not with RNA_ECO_. **(C)** CFU of ECO after co-incubation of PMN treated or not with RNA_PAE_. **(D)** The percentage of phagocytosis was evaluated by flow cytometry using ECO-GFP. PMN were incubated with ECO-GFP for 1 h (MOI 1) prior stimulation of 1 µg/mL RNA (RNA_PAE_ or RNA_ECO_). The percentage of association and adhesion of GFP-bacteria to PMN was measured at 37°C and 4°C, respectively. Percentage of phagocytosis was calculated by following subtraction: (%PMN-GFP^+^ 37°C) – (%PMN-GFP^+^ 4°C). Representative dot-plots of one independent experiment are depicted in the right. **(E)** Relative fluorescence (RF) of the mean fluorescence intensity (MFI) of GFP+ PMN of treated PMN relative to PMN without GFP+ ECO. A histogram plot of one representative experiment is depicted (right). All results are expressed as mean ± SEM (*p<0.05, **p<0.01).

According to the results showed above, we wondered whether the reduction on the ability of PMN to eliminate bacteria was specific for live PAE or whether elimination of live ECO could also be altered when PMN were pre-incubated with RNA_PAE_. Results depicted in Figure 6C show that RNA_PAE_ also reduced PMN ability to kill ECO.

In order to investigate the mechanisms by which RNA_PAE_ could be interfering with PMN bacteria killing, we evaluated phagocytosis by flow cytometry. PMN were treated or not with RNA_PAE_ and then were challenged with a GFP expressing ECO (ECO-GFP). PMN were incubated with ECO-GFP at 37 °C in order to detect bacterial association and at 4 °C to determine bacterial adhesion. The percentage of phagocytosis was calculated by subtracting these two conditions (37 °C – 4°C).

Figure 6D shows that the percentage of PMN with phagocytosed ECO-GFP was diminished when PMN were treated with RNA_PAE_. Also, when RNA_PAE_ was present, the amount of ECO-GFP *per* PMN on PMN-GFP^+^ gated cells was decreased (Figure 6E).

Altogether, these results indicate that RNA_PAE_ modulates PMN microbicidal activity favoring bacterial survival by decreasing phagocytosis.

### The integrity and size of RNA from *P. aeruginosa* are responsible of diminished microbicidal activity of PMN

To determine whether structural features of RNA_PAE_ were important for the decreased ability of PMN in bacterial elimination, we first digested RNA_PAE_ by adding RNAse A. Results on Figure 7A show that when RNA_PAE_ was degraded (RNA_PAE_+RNAse A), its ability to decrease PMN microbicidal ability was lost, as the number of surviving bacteria (CFU) was similar to that observed in unstimulated PMN. This result demonstrates that the capacity of RNA_PAE_ to decrease bacterial killing is dependent on RNA integrity. **Supplementary** figure 4A shows an agarose gel with isolated RNA and RNA+RNAse A (degraded RNA).

**Fig 7.**
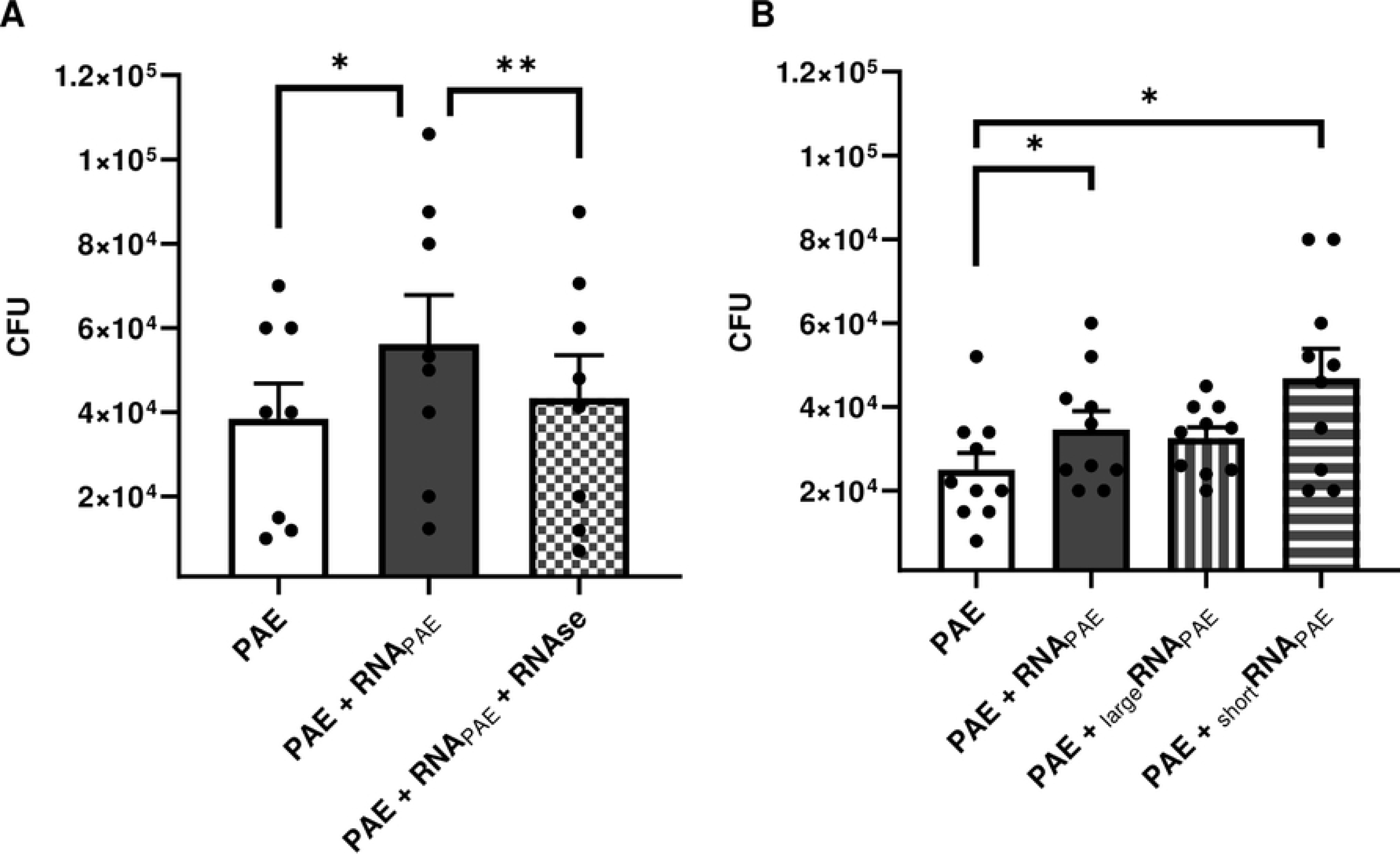
Bactericidal response on neutrophils (PMN) treated with degraded RNA_PAE_ or different RNA_PAE_ fractions that differ in size. **(A)** PMN were co-cultured with PAE for 1 h (MOI 1) in the presence or absence of 1 µg/mL RNA_PAE_ degraded or not with RNAse A, and the remaining viable colony forming units (CFU) were evaluated using TSA plates **(B)** PMN were co-cultured with PAE for 1 h (MOI 1) in the presence or absence 1 µg/mL of different size fraction of RNA_PAE_. RNA_PAE_ fractions are: _large_RNA (>150-200 bases) and _short_RNA (<150-200 bases). The remaining viable colony forming units (CFU) were evaluated using TSA plates after the incubation period. **(A and B)** Number of CFU. Each point represents an independent PMN donor. Results are expressed as mean ± SEM (*p<0.05, **p<0.01).

Additionally, we purified two different fractions of RNA according to its size: large RNA (_large_RNA) which is larger than >150-200 bp, and short RNA (_short_RNA) which contains RNA less than <150-200 bp. _large_RNA fractions contains large ribosomal RNA (23 S and 16 S subunits) and large mRNA, whereas _short_RNA contains small ribosomal RNA (5 S subunit), tRNA, small mRNA and microRNA. **Supplementary** figure 4B shows agarose gel with the different fractions of isolated RNA (total, large and short).

Killing assays were performed pre-stimulating PMN with 1 µg/ml of these different RNA fractions before challenging with live PAE. We found that no differences in CFU were found when _large_RNA was used compare to unstimulated PMN (Figure 7B). However, when _short_RNA from PAE was used, an increase in CFU was observed. This data suggest than only RNA smaller than 200 bp are responsible of reducing PMN ability to eliminate PAE.

In summary, here we described for the first time that RNA from PAE is able to diminish PMN responses against bacteria affecting PMN phagocytosis. Additionally, we found that short fragments of RNA are responsible for this effect and its integrity is necessary to reduce PMN responses.

## DISCUSSION

The severity and incidence of bacterial pneumonia, acquired mainly in health centers, and the problem that carries the lack of effective treatments against bacteria, which have a high resistance to antibiotics, imposes the need to study aspects related to the pathogenic mechanisms of these microorganisms. In recent years, prokaryotic RNAs (pRNA) have acquired great importance as modulators of the immune response (33). Here we included the RNA of two bacteria highly responsible for associated ventilator pneumonia*, E. coli* and *P. aeruginosa* (RNA_ECO_ and RNA_PAE_) (34,35).

Since the interaction between PMN and cells from the lung barrier is crucial in the resolution of infections, we did not only focused on the direct effects of RNA on PMN response and bacterial clearance but decided to include lung epithelial and microvascular endothelial cells in the study in order to have a global scenario that better represents an infectious focus.

In concordance with previous results (25,26), we found that RNA_ECO_ effectively acts as a stimulator not only for PMN and endothelial cells (HMEC-1) but also showed to be stimulatory for lung epithelial cells (Calu-6). In this regard, RNA_ECO_ induced Calu-6 and HMEC-1 activation observed as an increase in ICAM-1 expression and an increased secretion of IL-6 and IL-8. Moreover, CM from RNA_ECO_-treated cells were able to activate PMN. These results expand our understanding of the ability of RNA_ECO_ to induce a stimulatory response of PMN in a pulmonary context, where the influence of RNA_ECO_ on different steps or PMN activation is decisive, promoting the arrival of PMN at the site of infection by stimulating chemotaxis and adhesion. The induction of pro-inflammatory responses described here for RNA_ECO_ is in line with what was previously reported for RNAs of different microorganisms (both single- and double-strand RNA) on human or murine immune cells, such as dendritic cells and monocyte/macrophages (17,36–39)

Surprisingly, in the case of the RNA_PAE_, the scenario was completely different. RNA_PAE_ failed to induce any of the responses described for RNA_ECO_. Namely, no cell activation or cytokine secretion was induced in PMN, endothelium, or lung epithelial cells, and no PMN chemotaxis was observed when RNA_PAE_ was used as a stimulus. Moreover, when we determined the effect of RNA_PAE_ on PMN in the context of infection with live bacteria, we discovered that the absence of effects was not due to a simple deficiency of RNA_PAE_ for stimulating cells, but rather that RNA_PAE_ significantly decreased the ability of PMN to respond to live bacteria. In this sense, PMN activation induced by live PAE, observed as an increase in CD11b expression and PMN chemotaxis, was significantly reduced when RNA_PAE_ was added. More importantly, the ability of PMN to eliminate bacteria in the presence of the RNA_PAE_ was also decreased as a significant increment in the CFU of PAE was found. This reveals an immune evasion mechanism executed by the presence of RNA_PAE_ that favors the survival of PAE. Moreover, we also demonstrated that RNA_PAE_ not only reduced the PMN killing capacity against live PAE but also against live ECO, indicating that the reduction in bacterial clearance by PMN induced by RNA_PAE_ is not specific to PAE.

Previous evidence demonstrating that RNA may be used by specific pathogens to down-modulate the immune system is available in the literature. In this regard, Barrionuevo *et al*. have demonstrated that *Brucella abortus* RNA inhibited the MHC-I and MHC-II expression on the surface of macrophages (40,41). Consequently, the bacteria avoid the recognition by CD8+ and CD4+ T cells and evade the surveillance of the immune system. Moreover, Saha *et al.* documented that ssRNA from the Hepatitis C virus induces the differentiation of monocytes into macrophages with high expression of M2 surface markers and anti-inflammatory cytokine production (42). Our work supports the idea of bacterial RNA as a possible strategy used by some pathogens to evade the immune response and reports for the first time the influence of bacterial RNA in the context of a simulated (*in vitro)* bacterial infection. Previous studies describing the ability of RNA_PAE_ to modulate the immune response are scarce. To our knowledge, there is only one work of Koeppen *et al*. describing that a fragment of a *P. aeruginosa* methionine tRNA, packed on outer membrane vesicles (OMVs), was able to reduce IL-8 secretion induced by LPS in cultured primary human airway epithelial cells as well as cytokine secretion and PMN infiltration in mouse lung (21). We did not analyze RNA contained in extracellular vesicles but instead used purified RNA. It is believed that RNA may be released as a consequence of bacterial killing. Therefore, in light of our findings, in a physiopathogenic scenario we hypothesize that during a PAE infection, migration and activation of PMN may initially occur, but after a first round of bacterial killing, the release of RNA by this dead bacterium may attenuate further PMN migration and/or activation to give an advantage to the pathogen.

In an attempt to delve into the reasons why RNA_PAE_ attenuated the bactericidal response of PMN, we investigated whether phagocytosis of bacteria could be affected, and found that RNA_PAE_ reduced both the percentage of PMN with internalized ECO-GFP and the amount of internalized ECO *per* cell, while RNA_ECO_ did not alter these parameters. This novel finding could explain the increase in bacterial survival induced by RNA_PAE_.

Finally, our understanding of the modulatory ability of RNA_PAE_ to alter PMN functions was completed by revealing two additional characteristics of RNA_PAE_ necessary to reduce PMN bactericidal response. First, the integrity of RNA_PAE_ was an essential requirement, given that its degradation reversed the survival advantage of PAE compared to non-degraded RNA. Other authors have demonstrated that the integrity of the RNA is essential to regulate certain immunomodulatory responses of RNAs. In this sense, Rodrigues *et al*. have described that degradation of RNA from ECO negatively impacted its ability to stimulate PMN bactericidal mechanism (ROS generation and NETosis) but did not influence PMN activation overall, since CD11b up-regulation and increased chemotaxis were still observed when the RNA was degraded (25). This raises the concept that the integrity of bacterial RNA would be necessary to modulate some responses in PMN but not all. Second, by isolating the RNA fractions according to their length, we have also demonstrated that short fractions corresponding to <200bp were responsible for attenuating the killing capacity of PMN. It should be noted that Koeppen *et al.*also found that only short-sized RNA derived from PAÉs methionine tRNA suppressed the pro-inflammatory responses in human airway epithelial cells (21). Given that at the moment we cannot assure whether conformational structures or specific sequences are the determinants responsible for our findings, future experiments will be designed to shed light on these aspects.

Considering the results of the present study, we propose that once PAE colonizes the lung, it could activate adjacent epithelial and endothelial cells, inducing the secretion of cytokines and chemokines. This would cause the activation and migration of PMN to the infectious foci. These PMN will probably become further activated by bacteria and will display their microbicidal mechanisms. As bacterial death and lysis occur, the accumulation of extracellular RNA_PAE_ in the focus would lead to a decrease in bacterial phagocytosis and PAE killing by PMN, and also a reduction in new waves of PMN activation and arrival to lung. This would clearly guarantee an advantage for the persistence of PAE (Figure 8).

**Fig 8.**
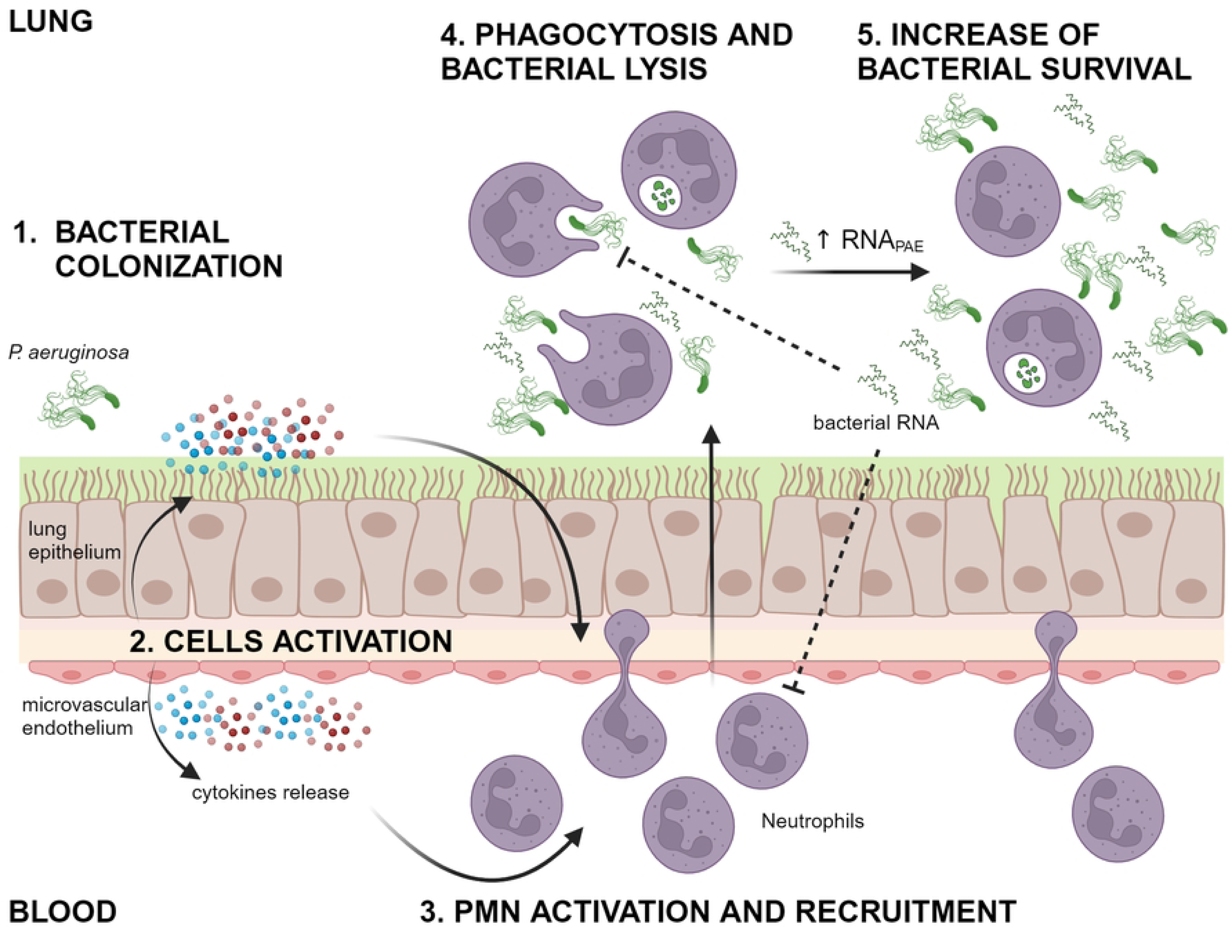
RNA from *P. aeruginosa* reduces PAE elimination by PMN in lung. **1.** PAE infects lung and initiates colonization**. 2.** Epithelial lung cell and microvascular endothelium senses PAMPs from PAE; cells turn activated and release pro inflammatory cytokines. **3.** Release of cytokines from epithelial lung cells and endothelium induces PMN activation and recruitment to the site of infection. **4.** PMN elimination and PAE, lysis causes an increment in free RNA from PAE. **5.** RNA_PAE_ diminishes phagocytosis by PMN and further activation and arrival of blood PMN, favoring bacterial survival and persistence. Created with Biorender.com

In summary, we have revealed a new immunosuppressive role of short RNA from PAE on PMN that gives a survival advantage to the bacteria by modulating PMN activation and killing capacity.

## MATERIALS AND METHODS

### Ethics Statement

Human normal samples were obtained from voluntary donors. This study was performed according to institutional guidelines (National Academy of Medicine, Buenos Aires, Argentina) and received the approval of the institutional ethics committee and written informed consent was provided by all the subjects.

### Bacterial Cultures

Experiments were performed using *Pseudomonas aeruginosa* (ATCC® 27853™) and *Escherichia coli* (ATCC® 25922™). Bacteria were grown in Tryptic Soy Broth (TSB) (Britania) for 18 h at 37°C. After that, 100 µL of the culture was added into 10 mL of fresh TSB, and grown for another 4 h with agitation, until the organism reached log phase. Bacteria was pelleted by centrifugation at 6000 g for 10 min, washed twice in phosphate buffered saline (PBS) 1x, and resuspended at the desired concentration. Bacteria concentration was determined by measuring O.D. at 600 nm, and adjusting to 0.09 absorbance units, that is equivalent to 1.10^8^ colonyforming units (CFU)/mL; CFU concentration was confirmed by counting on Tryptic Soy Agar (TSA).

### RNA Isolation

Bacterial (10.10^8^ CFU/mL) pellets were homogenized in RNAzol-RT (MRC Gene, Cincinnati, OH) to obtain prokaryotic RNA (pRNA). RNA was isolated from the resulting supernatant by alcohol precipitation, followed by washing and solubilization. Extraction controls without any biological sample were used for all untreated cells (control) to determine whether traces of phenol and thiocyanate compounds, which could remain after the extraction of RNA, had any effect on cell (PMN and cells lines) functionality. pRNA concentration was determined using a DeNovix DS11 Spectrophotometer (DeNovix Inc., Wilmington, DE). All samples had a 260/280 ratio between 1.8 and 2.1, and a 260/230 ratio between 1.8 and 2.3, indicating high purified RNA without phenol contamination. In some experiments, pRNA was treated or not with Ribonuclease A (RNase A; Merck Millipore, Darmstadt, Germany). For total degradation, a final concentration of 5 µg/mL was used, and the samples were incubated at 37°C for 20 min. After RNase A incubation, total digestion of pRNA was checked by agarose gel electrophoresis as described below. All experiments were performed in a final pRNA concentration of 1 µg/mL unless otherwise specified. Isolation of RNA fractions (large RNA and short RNA) were performed according to manufacturer’s instruction.

### Agarose Gel Electrophoresis

RNA fragments were separated using 2% agarose gel electrophoresis. 10 mg/mL of ethidium bromide was added to visualize the samples. Electrophoresis was performed at 60 V for 45 min. The RNA was detected using UV light and its size was determined using a standard 50 bp Plus DNA ladder (TransGen Biotech Co., Ltd, Beijing, China).

### Blood Samples

Blood samples were obtained from healthy volunteer donors who had not taken any medication for at least 10 days before the day of sampling. Blood was obtained by venipuncture of the forearm vein and was drawn directly into citrated plastic tubes.

### Polymorphonuclear Neutrophil (PMN) Isolation

Neutrophils were isolated by Ficoll-Hypaque gradient centrifugation (Ficoll Pharmacia, Uppsala; Hypaque, Wintthrop Products, Buenos Aires, Argentina) and dextran sedimentation (43). Contaminating erythrocytes were removed by hypotonic lysis. After washing, the cells (96% neutrophils on May Grünwald/Giemsa-stained Cyto-preps) were suspended in RPMI 1,640 supplemented with 2 % heat-inactivated fetal bovine serum (FBS) and used immediately after.

### Cell lines cultures

A cell line derived from human lung epithelium (Calu-6) was used. Cells were maintained in MCDB 131 (Gibco, Thermo Fisher Scientific, Waltham, MA, USA) supplemented with 10% fetal bovine serum (FBS) (Gibco, Thermo Fisher Scientific, Waltham, MA, USA), penicillin (100 U/mL), and streptomycin (100 mg/mL) (Sigma-Aldrich, MI, USA), and incubated at 37°C in 5% CO_2_. A human microvascular endothelial cell line (HMEC-1) was used. Cells were maintained in MCDB 131 (Gibco, Thermo Fisher Scientific, Waltham, MA, USA) supplemented with 10% fetal bovine serum (FBS) (Gibco, Thermo Fisher Scientific, Waltham, MA, USA), epithelial growth factor (10 ng/mL), hydrocortisone (1 μg/ml), L-glutamine (10 mM), penicillin (100 U/mL), and streptomycin (100 mg/mL) (Sigma-Aldrich, MI, USA), and incubated at 37°C in 5% CO_2_.

### Experimental Design

Calu-6 or HMEC-1 were seeded in 24-well plates and left until 90% of confluence was achieved. Cells were treated with RNA extraction control (Control) or isolated pRNA 1 µg/mL or 5 µg/mL depending on the experiment during 4 h (for cell conditioned media assay) or 18 h for cytokine determination and CD54 measurement.

5.10^5^ Isolated PMN were treated with RNA extraction control (Control) or isolated pRNA 1 µg/mL or 5 µg/mL, depending on the experiment, 30 min for CD11b determination, 4 h for cytokine secretion or 20 min before challenged with live bacteria.

### Cell conditioned medium

Calu-6 or HMEC-1 were seeded in 24-well plates and left until 90% of confluence was achieved. Cells were treated with 1 µg/mL of pRNA, unless otherwise specified, for 4 h, and then cells were gently washed three times with PBS 1X. Then, complete fresh medium was added, and cells were left for an additional period of 18 h. After this, medium was collected (conditioned media, CM) and used immediately for experiments.

### Flow Cytometry Studies

Cells were analyzed on a FACScalibur flow cytometer (BD Biosciences) and data were processed with the FlowJo 7.6.2 or vX.0.7 applications (FlowJo, LLC). For ICAM-1 (CD54) measurement, cell lines were treated, trypsinized, washed and stained with a specific anti-CD54 human antibody conjugated with phycoerythrin (PE, eBioscience,Thermo Fisher Scientific) at 4°C for 30 min. For CD11b measurement, 2,5.10^5^ PMN were incubated with different treatments and then stained for 30 min with a specific mouse anti-human CD11b antibody conjugated with PE (Dako, Santa Clara, CA, USA) at 4°C for 30 min. Debris were excluded by FSC-SSC and CD54 or CD11b increase expression was analyzed within gated-viable cells. Mean fluorescence intensity (MFI) was determined on 20.000 events. Isotype matched control immunostaining was performed in parallel.

### Viability assay

For viability assays, cells were treated with pRNA 5 µg/mL for 18 h in cell lines and 4 h in PMN. Cells treated with 2% paraformaldehyde (PFA) were included as a positive control in cell lines and phorbol ester, phorbol-12-myristate-13-acetate (PMA, 100 nM) (Thermo Fisher Scientific, Waltham, MA, USA) as a positive control for PMN death. After treatment, cells were incubated with 7-AAD (7-aminoactinomicine, BD Biosciences) for 10 min on ice and in the dark. All 7-AAD negative cells were considered viable.

### Cytokine determination

IL-6 and IL-8 were determined in cell lines supernatants stimulated for 18 hours with pRNA or PMN supernatants after 4 h. Supernatants were centrifuged to eliminate cells and debris and kept at -20°C until interleukin determination. Release of IL-6 and IL-8 were measured using commercial ELISA kits (Biolegend, San Diego, CA, USA) according to manufacturer’s instructions. Results were expressed in pg/mL and were extrapolated from serial dilutions.

### PMN adhesion to cells

PMN adhesion to cells was determine measuring the activity of alkaline phosphatase (ALP) from adherent PMN as previously described (44). Briefly, purified PMN were seeded onto treated HMEC-1 or Calu-6 in a ratio of 20:1 and cultured for 90 min. Then, non-adherent PMN were removed by vigorous washing with PBS. ALP activity of remaining adherent PMN was performed by measuring the conversion of p-nitrophenylphosphate substrate to nitrophenol. For this, adhered cells were incubated for 2 h at 37°C with 100 µL of 1% p-nitrophenylphosphate (2 mg/mL, Sigma-Aldrich) in AMP buffer (0.1 M, pH = 10) with 0.2 % SDS and 1 mM Cl_2_Mg. Then, 50 µL of stop solution (2 N NaOH) was added and optical density was measured at 405 nm in an AsysUVM340 Microplate Reader (Biochrom Ltd., Holliston, MA, USA). 1 Unit of ALP was defined as the amount of enzyme that produces 1 µmol of nitrophenol/min.

### Chemotaxis assay

Chemotaxis was performed using a modification of the Boyden chamber technique (45). Briefly, cell suspension (50 µL) containing 2.10^6^ PMN/ml in RPMI with 2% FBS, was placed in the top wells of a 48-well micro-chemotaxis chamber. A PVP-free polycarbonate membrane (3 µm pore size; Neuro Probe Inc. Gaithersburg MD, USA) separated cells from lower wells containing either RPMI or the stimulus (pRNA or CM). All samples were performed by triplicate. The chamber was incubated for 30 min at 37°C in a 5% CO_2_ humidified atmosphere. After incubation, the membrane was stained with TINCION-15 (Biopur SRL, Rosario, Argentina), and the number of PMN on the undersurface of the membrane was counted in five random high-power fields (HPF) × 1000 for each triplicate sample.

### Phagocytosis

Bacterial phagocytosis was determined as before (46). Briefly, PMN (1.10^6^) were incubated with 1.10^6^ CFU of GFP-ECO for 1 h at 37 °C (total bacteria-PMN interaction) or 4 °C (for bacterial adhesion) in 5 % CO_2_. After incubation, GFP^+^ PMN were evaluated by flow cytometry and expressed as the percentage of phagocytosis which resulted from subtracting PMN associated to GFP-bacteria (37 °C) minus the percentage of PMN with attached bacteria (4 °C).

### Bactericidal Activity Assays

Bacterial killing by PMN was determined using PMN (1.10^6^) previously stimulated for 20 min with pRNA (1 µg/mL) in a 24-well tissue culture plate, and then challenged with 1.10^6^ CFU of bacteria. One hour after incubation PMN were lysed with H_2_O, and this solution was plated on TSA in serial dilutions. CFU were enumerated the following day.

### Statistical Analysis

Results were expressed as the mean ± SEM. Statistical analysis of the data was performed using the Mann-Whitney or T-test for two group comparisons according having or not a Gaussian distribution. Comparisons among groups were made by analysis of variance (ANOVA), applying Bonferroni or Kruskal-Wallis according having or not a Gaussian distribution. p<0.05 were considered significant

## AUTHOR CONTRIBUTIONS

**Conceptualization:** José R Pittaluga, Federico Birnberg-Weiss, Gabriela C Fernández, Verónica I Landoni.

**Data curation:** José R Pittaluga, Verónica I Landoni.

**Formal analysis:** José R Pittaluga, Verónica I Landoni.

**Funding acquisition:** Paula Barrionuevo, Gabriela C Fernández, Verónica I Landoni.

**Investigation:** José R Pittaluga, Federico Birnberg-Weiss, Joselyn Castro, Agustina Serafino, Luis A Castillo, Daiana Martire-Greco, Paula Barrionuevo, Gabriela C Fernández, Verónica I Landoni.

**Methodology:** José R Pittaluga, Federico Birnberg-Weiss, Gabriela C Fernández, Verónica I Landoni.

**Resources:** Luis A Castillo, Paula Barrionuevo, Gabriela C Fernández, Verónica I Landoni.

**Supervision:** Gabriela C Fernández, Verónica I Landoni.

**Visualization:** José R Pittaluga

**Writing – original draft:** José R Pittaluga, Gabriela C Fernández, Verónica I Landoni.

**Writing – review & editing:** José R Pittaluga, Federico Birnberg-Weiss, Joselyn Castro, Gabriela C Fernández, Verónica I Landoni.

## CAPTIONS OF SUPPORTING INFORMATION

**S1 Fig.RNA from P. aeruginosa failed to activate epithelial lung or endothelial cells even at higher concentrations.** Calu-6 and HMEC-1 cell lines were unstimulated (CONTROL) or stimulated with 5 µg/mL RNA (RNAPAE or RNAECO) for 18 h. ICAM-1 expression on cell surface by flow cytometry,expressed as the relative fluorescence (RF) of the mean fluorescence intensity (MFI) of treated samples relative to isotype control.Histogram plots from one representative experiment aredepictedbelow.(A) Calu-6 ICAM-1 expression, (B) HMEC-1 ICAM-1 expression. Results are expressed as mean ± SEM (**p<0.01, ***p<0.001, ****p<0.0001).

**S2 Fig. Cell viability is not affected after incubation with a high concentration of pRNA. Calu-6 and HMEC-1 cell lines were unstimulated (CONTROL) or stimulated with 5 µg/mL RNA (RNAPAE or RNAECO) for 18 h.** 7-AAD staining was measured in cells by flow cytometry, expressed as the percentage of 7-AAD+ cells. PFA (2 %) was used as a positive control for 7-AAD incorporation. (A) Percentage of 7-AAD+ Calu-6 cells. (B) Percentage of 7-AAD+ HMEC-1 cells. (C) PMN were unstimulated (CONTROL) or stimulated with 5 µg/mL RNA for 4 h to determine 7-AAD staining by flow cytometry. PMA (100 nM) was used as a positive control for cell death. Results were expressed as the percentage of 7-AAD+ PMN. Histogram plots from one representative experiment are presented (right). Results were expressed as mean ± SEM (****p<0.0001).

**S3 Fig. RNA from P. aeruginosa failed to activate neutrophils even at higher concentrations.** PMN were unstimulated (CONTROL) or stimulated with 5 µg/mL RNA (RNAPAE or RNAECO) for 30 min to determine CD11b expression by flow cytometry. (A) CD11b expression, expressed as the relative fluorescence (RF) of the mean fluorescence intensity (MFI) of treated samples relative to isotype control. Histogram plots from one representative experiment is presented (right).Secretion of IL-8 was measured by ELISA in the supernatant of PMN after 3 hours of stimulation. (B) IL-8 concentration. Results are expressed as mean ± SEM (**p<0.01, ***p<0.001, ****p<0.0001).

**S4 Fig. Representative agarose gel electrophoresis stained with ethidium bromide. M-Marker: 50 bp Plus DNA ladder.** (A) Total RNA from PAE and RNA treatment with RNAse. (B) RNA fractions isolation: Total RNA, large RNA and short RNA from PAE. White lines indicate different lanes of the same gel

